# The relationships between structural organization, material properties, and loading conditions and the risk of fracture and fracture location in the femur

**DOI:** 10.1101/2021.07.26.453913

**Authors:** Todd L. Bredbenner

## Abstract

Increased risk of skeletal fractures due to bone mass loss is a major public health problem resulting in significant morbidity and mortality, particularly in the case of hip fractures. Current clinical methods based on two-dimensional measures of bone mineral density (areal BMD or aBMD) are often unable to identify individuals at risk of fracture. The underlying hypothesis of this study was that combinations of femur structural traits are different for those femurs that suffer a fragility fracture within the proximal region of the femur and those that sustain a fracture in either the subtrochanteric or midshaft region of the femur, resulting in an “atypical femur fracture”. Accordingly, the objective of this study was to determine the effects of varying combinations of structural traits, material properties, and loading conditions on femur stress response and the location of stress response variation using a validated parametric finite element model. Statistical shape and trait modelling of the femur was used to describe variability in the structural organization of a set of femurs in an efficient manner and the resulting description of structural variability was exploited to investigate how different mechanisms of fracture might occur, whether in the proximal region or in the subtrochanteric and midshaft region. In combination with parameters describing loading condition and material property variation, variation in structural organization is associated with regional increases in maximum principal stress and the percentage of bone expected to damage, and these increases are likely associated with increased fracture risk. The results of this study indicate that there are multiple pathways and combinations of descriptor variation that may result in increased fracture risk and that these pathways can lead to fracture in any region of the femur under both overload conditions, such as with sideways fall loading, and stance loading, which due to the repetitive nature may lead to the accumulation of fatigue damage within the bone and further impair bone condition and increased susceptibility to fracture.

## INTRODUCTION

Osteoporosis is an enormous and growing public health problem in the aging population with associated health care costs of nearly $18 billion in the U.S. [1]. It is estimated that 40%-46% of women over 50 and 13%-22% of men over 50 will suffer an osteoporosis-related fracture [2]. With the number of persons over 60 projected to more than double by 2050, the aging of the general population will lead to a substantial rise in the at-risk population for fractures and the overall costs associated with osteoporotic fractures could double or triple in the coming decades [3, 4].

Typically, fracture risk is assessed in the clinic using bone mineral density (BMD), a measure of the amount of bone mineral in a region of interest. BMD is routinely measured using dual energy x-ray absorptiometry (DXA) to determine areal BMD (aBMD), a two-dimensional averaged description of bone mineral content. However, it is evident that aBMD alone does not explain the likelihood of atypical femur fracture and that bone quantity measures need to be complemented with more comprehensive measures of bone structure to substantially increase the accuracy of fracture risk assessment [5]. Bone quality (e.g., arrangement and composition of bone) is a critical component of bone fracture resistance, yet little is known about the interaction of the bone quality traits with overall geometry and distributions of BMD and material properties that in combination are responsible for the integrity of skeletal structures, such as the femur, under the range of potential loading conditions.

The current research paradigm in fracture biomechanics focuses strongly on BMD and its contribution to bone fragility and is dominated by investigations of the role of one or a small group of closely related traits. In fact, the structural integrity of bone in any mechanical loading environment is an integrative function of a multitude of complex and interrelated characteristics of bone at the various levels of structural organization. Consequently, a great deal of osteoporosis fracture risk is independent of BMD [6, 7].

Statistical shape modeling methods have been used to describe variability in the morphology of a population of anatomical structures in terms of a random field representation [8-11]. Statistical shape models capture the variability of biological structures by projecting a high dimensional representation of the structure onto a lower dimensional subspace of possible shapes constructed from a population of training shapes. Additionally, the framework of statistical shape modeling allows for description of variation in spatial distributions of traits, such as the distribution of bone mineral density (BMD) within a bone [12, 13]. A modeling approach combining statistical shape and trait modeling with finite element modeling allows investigation of the response to loading in specific individuals, as well as over the full range of morphological and trait variability described within the model set. Statistical shape and trait modeling and variants have been used to investigate differences in the size, shape, and bone mineral density distributions of bones that suffer a fracture versus that that do not suffer a fracture within some follow-up period [12].

In recent years, the topic of “atypical femur fractures” has been of concern in relation to osteoporosis and long-term osteoporosis treatment. As opposed to the proximal femur fractures generally associated with osteoporosis, atypical femur fractures occur in the subtrochanteric or diaphyseal region, which is generally the strongest part of the femur [14-16]. In addition to the possible, but not clearly established, link between atypical fractures and long-term bisphosphonate treatment [17], there are a number of femur traits, particularly with regards to geometry, that have been investigated for possible relevance to both “typical” proximal femur fractures and atypical femur fractures, including proximal and overall femur geometry, cortical thickness, regional variation in trabecular bone density and material property variation related to changes in bone turnover [18-31]. However, these traits are typically investigated for associations with fracture risk without consideration of the inherent interaction between the traits that is structurally responsible for the loading response of the femur.

This study was based on the underlying hypothesis that combinations of femur structural traits are different for those femurs that suffer a fracture in the subtrochanteric or diaphyseal region, as opposed to a fracture in the femoral neck or intertrochanteric regions. The objective of this study was to determine the effects of varying combinations of structural traits, material properties, and loading conditions on femur stress response predictions and the location of increased stress using a validated parametric finite element model.

## METHODS

### Experimental Testing of Intact Femurs

A set of fifteen right cadaver femurs was obtained (Willed Body Program, UT Southwestern Medical Center, Dallas, TX). The femurs were stripped of soft tissue and the distal end was potted to the level of the condylar flare using a fast-curing acrylic resin (Clarocit, Struers, Inc., Cleveland, OH). The femurs were positioned to simulate sideways-fall loading conditions with an internal rotation angle of 15º and an angle of 10º between the horizontal and the mechanical axis of the femur [32]. The greater trochanter was positioned on a smooth platen above a load cell (22 kN capacity, Interface Inc., Scottsdale, AZ) in a custom-built servo-hydraulic load frame and the distal femur was mounted to allow only rotation about a horizontal axis perpendicular to the femoral shaft. Load was applied to the femoral head under displacement control at a rate of 100 mm/sec. until the femur failed. Greater trochanter load and crosshead displacement were recorded throughout loading.

### Image Processing

Computed tomography (CT) image data for forty-seven intact cadaver specimens (39 males, 64.3 ± 8.6 years; 8 females, 67.1 ± 12.7 years) was obtained (University of Virginia Center for Applied Biomechanics, Charlottesville, VA). The femurs were scanned using a computed tomography (CT) system (Optima CT660, GE Healthcare, Waukesha, WI) and reconstructed with 0.977 × 0.977 × 0.625 mm^3^ voxels. QCT data was filtered using a sequence of median and anisotropic diffusion filters to reduce data noise. Filtered data was semi-automatically segmented to extract right femur data from the CT image data (MATLAB R2017a, The Mathworks, Inc., Natick, MA; Seg3D, The Center for Integrative Biomedical Computing, University of Utah, Salt Lake City, UT). Except as noted, MATLAB was used for the remainder of data processing. Using a marching cubes algorithm, watertight triangulated surfaces were generated to describe the outer cortical boundary of each femur by computing the isosurface geometry for the segmented data region and smoothing the resulting surface to remove any stair-stepping effects due to image resolution. Femur surfaces were resampled, resulting in approximately 10,000 faces for each triangulated surface.

All femur surfaces were positioned so that the posterior-most aspects of the femoral condyles were tangential to the same plane defined by the CT scanner axes. A femur surface was arbitrarily selected as the template and additionally aligned with the vertical CT axis. Remaining surfaces were aligned to the template surface.

Vertices from the template surface were mapped onto the remaining femur surfaces. Using a coherent point drift algorithm followed by a non-rigid iterative closest point algorithm, vertices were repositioned such that all vertices were positioned at corresponding anatomic locations for all femur surfaces [33, 34]. Thus, the resulting outer femur surfaces were defined by the same surface definition due to vertex correspondence across the set of femurs.

An optimized full-width half-maximum algorithm was used to estimate cortical thickness from the CT data in order to determine the transition from cortical bone to trabecular bone and marrow space [20, 35]. CT data was processed along with the corresponding outer surface definition for each femur to define an inner cortical surface that was also corresponding across the set of femurs and shared the same triangulated surface definition as the outer surface.

### Development of Finite Element Models of Individual Femurs

A volumetric tetrahedral mesh consisting of approximately 28,000 elements (7,000 nodes) was defined for the template outer surface and refined to improve mesh quality (Tetgen, Weierstrass Institute for Applied Analysis and Stochastics, Berlin, Germany). The resulting femur mesh was warped using a nonlinear elastic method to match the remaining periosteal surfaces using displacement vectors calculated between corresponding surface vertices on the template mesh and each individual periosteal surface, resulting in a set of forty-seven corresponding femur mesh models that were aligned to each other (LS-DYNA R10.10, Livermore Software Technology Corporation, Livermore, CA).

Individual volumetric models were superimposed on the CT data for the same individual and CT image intensity was determined at the spatial location of each node. The image intensity distribution for each femur was converted to Hounsfield units and then to BMD using a regression based on the graded K_2_HPO_4_ densities within the solid water equivalent K_2_HPO_4_ density calibration phantom and image intensity data within corresponding regions of interest. The distribution of equivalent K_2_HPO_4_ concentration was converted to a wet apparent bone density distribution for each individual femur [36]. This process resulted in a set of forty-seven femur geometry and density distribution models where each model consisted of 49,000 variables (i.e., spatial location and wet apparent bone density at each femur mesh node, as well as spatial location of the vertices describing the inner cortical boundary).

For each femur, centroids were determined for each triangle in the outer and inner surfaces. The resulting centroid points were added as nodes in the tetrahedral mesh for each and the mesh was refined to incorporate the new nodes and to ensure tetrahedral element quality (Tetgen, Weierstrass Institute for Applied Analysis and Stochastics, Berlin, Germany). The resulting volumetric models had approximately 142,000 tetrahedral elements (∼27,000 nodes) and explicitly described the inner and outer cortical boundaries, as well as the trabecular and marrow regions of the femurs. Wet apparent bone density was determined at the centroid of each tetrahedral element by interpolating from the previous point-wise apparent density distribution for the same femur.

The range of wet apparent bone densities (0.0 – 2.0 mg/cm^3^) was divided into twenty equally-sized bins and the spatial distribution of apparent density was simplified to consist of twenty values. An isotropic elastic-plastic material model with asymmetric tension and compression yield behavior was employed to describe both trabecular and cortical bone in the femur models. Distributions of isotropic elastic moduli and compressive and tensile yield stresses were determined by applying empirical relationships to the distributions of wet apparent density. Elements with wet apparent density values of 1.0 mg/cm^3^ or below were assigned material properties based on trabecular bone density-property relationships [37, 38] and elements with wet apparent density values greater than 1.0 mg/cm^3^ were assigned material properties based on cortical bone relationships [39, 40]

The femur models were transformed to simulate sideways-fall loading conditions with an internal rotation angle of 15º and an angle of 10º between the horizontal and the mechanical axis of the femur [32]. Boundary conditions were applied to the models such that, below the condylar flare, translation of the femur was constrained and only rotation about a horizontal axis perpendicular to the femoral shaft was allowed. The greater trochanter was constrained from vertical translation, but the remaining degrees of freedom were unconstrained. Displacement was prescribed to a set of nodes on the medial side of the femoral head and the force on the greater trochanter was determined at 5% displacement of the femoral head relative to the unloaded position. Greater trochanter force was compared between the experimentally-loaded femurs and the individual femur models.

### Development of a Statistical Shape and Trait Model of the Femur

A statistical shape and trait model (SSTM) was generated to describe and investigate variability in femur geometry and BMD distribution within each femur and body weight. Joint point distribution models were constructed from the original forty-seven femur geometry (i.e., inner and outer cortical surfaces) and density distribution models and body weights for the individual. The original geometry, density, and body weight for each individual was described by a shape and trait parameter vector as

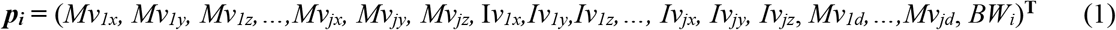

where *Mv*_*j(xyz)*_ are the three-dimensional coordinates of the nodes in the original tetrahedral femur model, *Iv*_*j(xyz)*_ are the three-dimensional coordinates of the vertices in the original inner surface, *Mv*_*jd*_ is the wet apparent bone density at each tetrahedral mesh node, *BW*_*i*_ is the body weight of the *i*^th^ individual, *j* =1,…, *J* = 7,000 nodes in the volumetric mesh, and *i = 1,…,n =47* denotes each individual in the set. The mean shape and trait description in the set of femurs and individuals was defined as

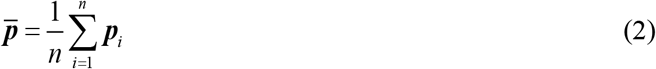

and the correlation between individual models in the set was given by the empirical covariance matrix

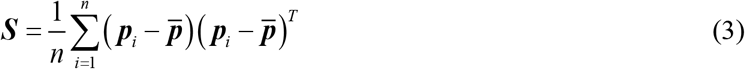

A principal components analysis of the covariance matrix, ***S***, results in a set of *k* = *n* - 1 eigenvalues (*λ*_*k*_) and eigenvectors (***q***_*k*_), which are the principal directions spanning a shape space centered at the mean, 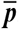. The proportion of the total variance described along each eigenvector is equal to its corresponding eigenvalue divided by the sum of all eigenvalues; eigenvectors corresponding to the largest eigenvalues describe the majority of the variance. Thus, the original tetrahedral mesh, inner surface, and wet apparent bone density for each femur and body weight for each individual in the set were described in terms of the average model and a weighted linear combination of uncorrelated principal shape modes as

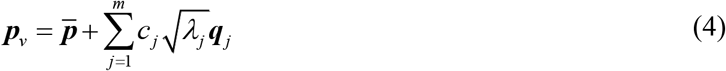

where ***p***_***v***_ is a vector containing coordinates for all node position and density information in the model, ***m*** is the number of eigenvalues, *λ*_*j*_, , and deviation from the average, 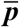, was determined as the sum of the products of a set of scalar weighting factors, ***c***_***j***_, and SSTM standard deviations, 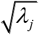, along the ***q***_***j***_(eigenvector) directions [12, 13].

Accordingly, the highly correlated three-dimensional geometry and density distribution variables are reduced into a relatively small set of uncorrelated and independent composite traits. All variability within the original set of femur models (originally described by over 49,000 variables) is now described by the weighting factors of 46 composite traits for each individual. Composite traits are new descriptive variables that, by definition, are linear combinations of the original descriptive variables and, furthermore, all geometry and trait information in the original models is retained in the new model descriptions.

### Development of a Parametric Finite Element Model of the Femur

The statistical shape and trait model of the femur was generated in a form directly applicable to finite element analysis and geometry and density variation in the finite element model was explicitly described by composite trait weighting factors. The effects of variation in femur geometry and density distribution were investigated by varying the weighting factors for a subset of composite traits describing 99% of the variability within the set of femurs. Composite trait weighting factors were defined as random variables with a mean, standard deviation, and distribution shape (Table 1). As with the individual femur models, a refined tetrahedral mesh consisting of approximately 142,000 elements with explicit inner and outer cortical boundaries could be created from the tetrahedral mesh, inner cortical surface, and density distribution associated with weighting factors determined by sampling the variable space.

**Table 1:** Random Variable Definitions

|  | Random Variables | Value | Distribution Type |
| --- | --- | --- | --- |
| FEMUR TRAITS | Composite Trait Weighting Factors | $0.0 \pm 0.31$ | Normal |
| LOADING | <u>Fall Loading</u> |  |  |
| CONDITIONS | Internal Rotation Angle | $15.0 \pm 4.5$ | Normal |
| | Shaft Angle | $10.0 \pm 4.02$ | Normal |
|  | <u>Stance Loading</u> |  |  |
| | Resultant Force Component (X) | $-0.31 \pm 0.08$ (-0.6 - -0.12) | Truncated Normal |
| | Resultant Force Component (Y) | $0.06 \pm 0.06$ (-0.06 - 0.27) | Truncated Normal |
| | Resultant Force Component (Z) | $-0.94 \pm 0.03$ (-0.99 - -0.79) | Truncated Normal |
| MATERIAL | Scale Factor (Trabecular, Modulus) | $1.0 \pm 1.5$ | Normal |
| PROPERTIES | Scale Factor (Cortical, Modulus) | $1.0 \pm 1.5$ | Normal |
| | Scale Factor (Trabecular, Yield Stress) | $1.0 \pm 1.5$ | Normal |
| | Scale Factor (Cortical, Yield Stress) | $1.0 \pm 1.5$ | Normal |

Material properties were assigned to trabecular and cortical bone elements, as before. However, random variables consisting of scale factors to modify the resulting spatial distributions of trabecular and cortical bone moduli and tensile and compressive yield strengths were defined to investigate the effects of variability in material property definition (Table 1).

One set of finite element models was created to model sideways fall loading, and another set was created to model single leg stance. For the fall loading models, random variables were defined to describe variation in the internal rotation and shaft angles [32, 41] (Table 1). Boundary conditions were defined as described for the individual femur models and displacement was identically prescribed to a set of nodes on the medial side of the femoral head. For the stance loading models, the direction of the femoral head resultant force was modeled using random variables [42] and distributed over a set of nodes on the superior surface of the femoral head (Table 1). The direction of a force vector simulating the abductor muscle force was defined in the opposite direction to that of the femoral head resultant [43-45] and applied to a set of nodes on the surface of the greater trochanter. The magnitude of the femoral head resultant was defined as 3.4 x body weight [42] and the magnitude of the abductor load was defined as 60% of the femoral head resultant magnitude [43, 44]. Body weight for each individual model was determined from the statistical shape and trait model following weighting factor definition.

A finite element model of the mean femur was created using the mean random variable values. Additional models to investigate the effects of variation in geometry, density distribution, material properties, and loading conditions were created by sampling the random variable distributions using a Latin Hypercube approach with 10 x (r +1) samples, where r = the number of random variables in the set of models for each loading condition (i.e., fall or stance) (NESSUS v9.8, Southwest Research Institute, San Antonio, TX). The sampled sets of random variables were used to define finite element models as previously described, resulting in a mean model and 180 variation models for sideways fall loading and a mean model and 190 variation models for stance loading. All models were subjected to appropriate loading conditions and solved to determine the stress distribution throughout the femur (LS-DYNA R10.10, Livermore Software Technology Corporation, Livermore, CA).

### Investigation of parameters related to increases in regional femur stress

The spatial distribution of principal stress was calculated for the upper half of each femur. In the case of fall loading, the ratio of maximum (tensile) principal stress to tensile yield strength was determined on an element-by-element basis. The upper half of each femur was divided into femoral neck, intertrochanteric, subtrochanteric, and midshaft regions and each spatial region was separated into cortical and trabecular bone regions. Regional stress response was quantified as the ratio of elements where the maximum principal stress was greater than the tensile yield strength (e.g., “yielded” elements) to the number of elements in each region. For stance loading, regional stress response was quantified as the maximum principal stress in each region, as the relatively low loads associated with stance did not lead to material yield.

A non-parametric Gaussian process model was fit to relate the distributions of input parameters (i.e., random variables) to femur stress response (NESSUS Response Surface Toolkit, Southwest Research Institute, San Antonio, TX). Global sensitivity analyses of the Gaussian process models were used to quantify the association of variation in femur stress distribution to femur structure, material properties, and loading variation for each loading type and each femur region (i.e., cortical and trabecular bone in each of the proximal femur and subtrochanteric femur, down to the mid-line of the femur) [46, 47]. Variance-based main effect indices directly quantified the association between each random variable and the variation in femur stress response. Total effect indices quantified the association between variation in random variables, including interactions with other random variables, with respect to variation in femur stress response.

Although an efficient means of describing variation in uncorrelated combinations of traits in femur geometry and density distribution, composite traits do not have explicit physical meaning, as they describe combinations of discrete structural descriptors, such as linear or angular measures of geometry and regional bone mineral density distribution measures. Therefore, descriptors of femur geometry and density distribution were determined for each region in all femurs. Femur curvature was determined in both the coronal and sagittal plane for the proximal and middle thirds of the femur and for the full femur [48]. Additional discrete measures of femur geometry, including neck axis length, neck length, neck diameter, neck axis-shaft angle, and overall femur length were determined for each femur [23, 24]. Finally, the mean wet apparent density for both cortical and trabecular bone compartments within each region was determined as a simple measure of bone density variation.

## RESULTS

### Experimental testing of intact femurs

The set of 15 cadaver right femurs was successfully tested in fall-type loading using a servo-hydraulic test frame. Fracture load was defined as the peak reaction load on the greater trochanter and the resulting fracture load distribution was 5,319.8 ± 2,240.6 N (2,942.9 – 9,785.6 N).

### Analysis of individual femur models

Computational solution of the 47 individual femur models resulted in a greater trochanter reaction load of 7,693.3 ± 1,815.9 N (4,390.0 – 11,340.0 N) at 5% deformation, which was taken as comparable to the experimental femur fracture load. An unpaired t-test determined that the mean femur fracture load for the individual femur models was significantly higher than that of the experimental femurs (7,693.3 N vs. 5,319.8 N, p-value < 0.005). Despite the higher mean femur load in the individual finite element models, the range of values was comparable to that of the experimental femurs.

### Variation in femur structure

Eleven composite trait weighting factors described 99% of the variability in femur structure (i.e., size, shape, and the spatial distributions of cortical thickness and bone mineral density), with 81% of the variability described by a single weighting factor (WF1) and the remaining 18% cumulatively described by 10 additional weighting factors (Figure 1).

**Figure 1:**
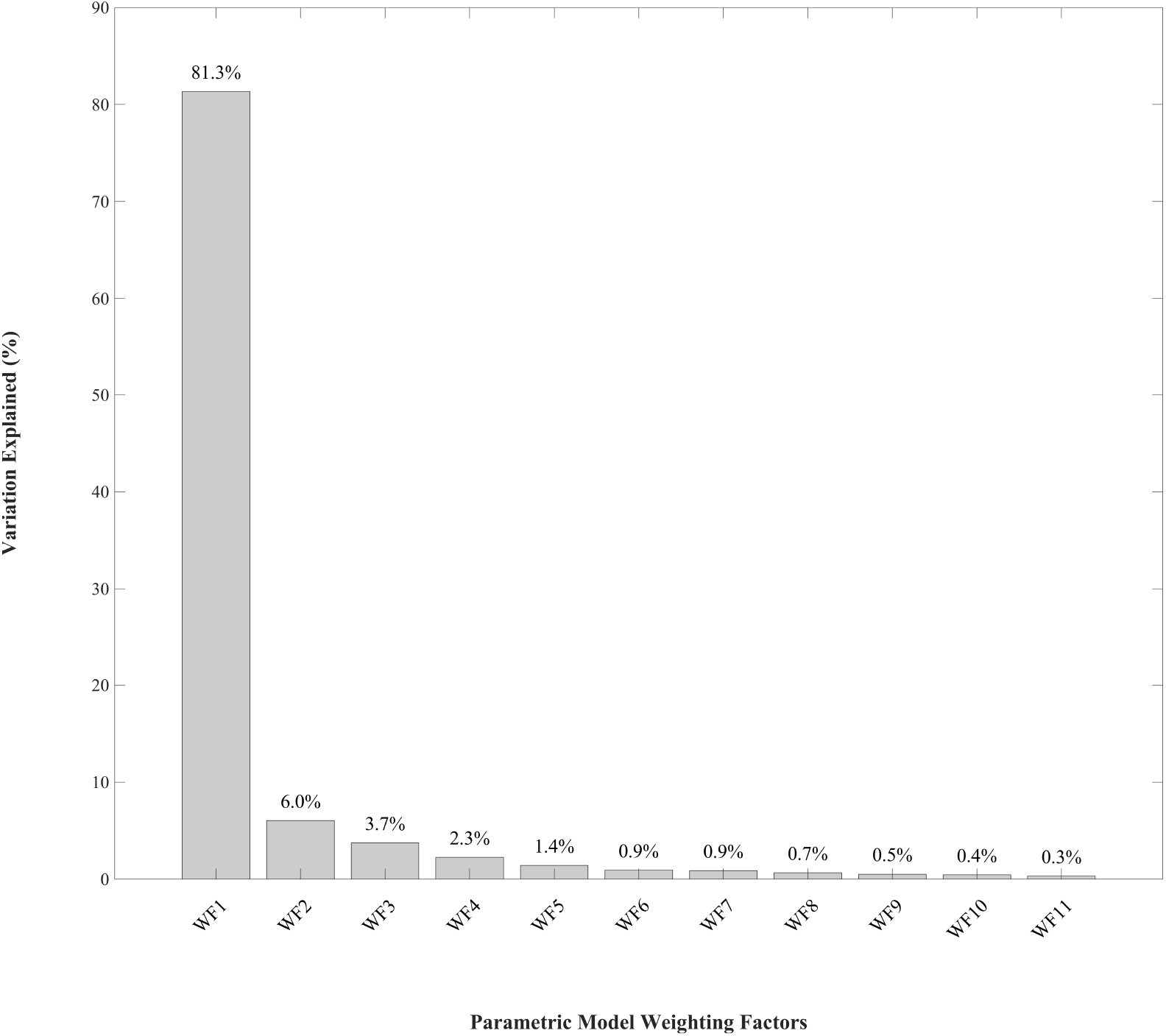
Variability explained by individual parametric model weighting factors

### Analysis of factors associated with the distribution of femur stress

Under simulated fall loading conditions, yield strength for trabecular bone was predominantly associated with variations in the percentage of yielded elements for both the cortical and trabecular compartments of the proximal femur (Figure 2). Trabecular bone stress in the proximal femur was also affected by the cortical bone modulus and variation in femur structure (WF4-WF9), although interaction between these factors and trabecular bone yield stress was associated with trabecular bone stress variation, rather than through the effects of these factors alone. Variation in subtrochanteric cortical bone stress was dependent on variation in cortical elastic modulus, yield strength for both cortical and trabecular bone, and a number of structural variables (Figure 3). Trabecular bone yield strength was primarily responsible for variation in subtrochanteric trabecular bone stress, although interaction with a number of femur structural variables contributed to stress variation (Figure 3). Loading variation had minimal effect on changes in stress distribution under fall loading conditions.

**Figure 2:**
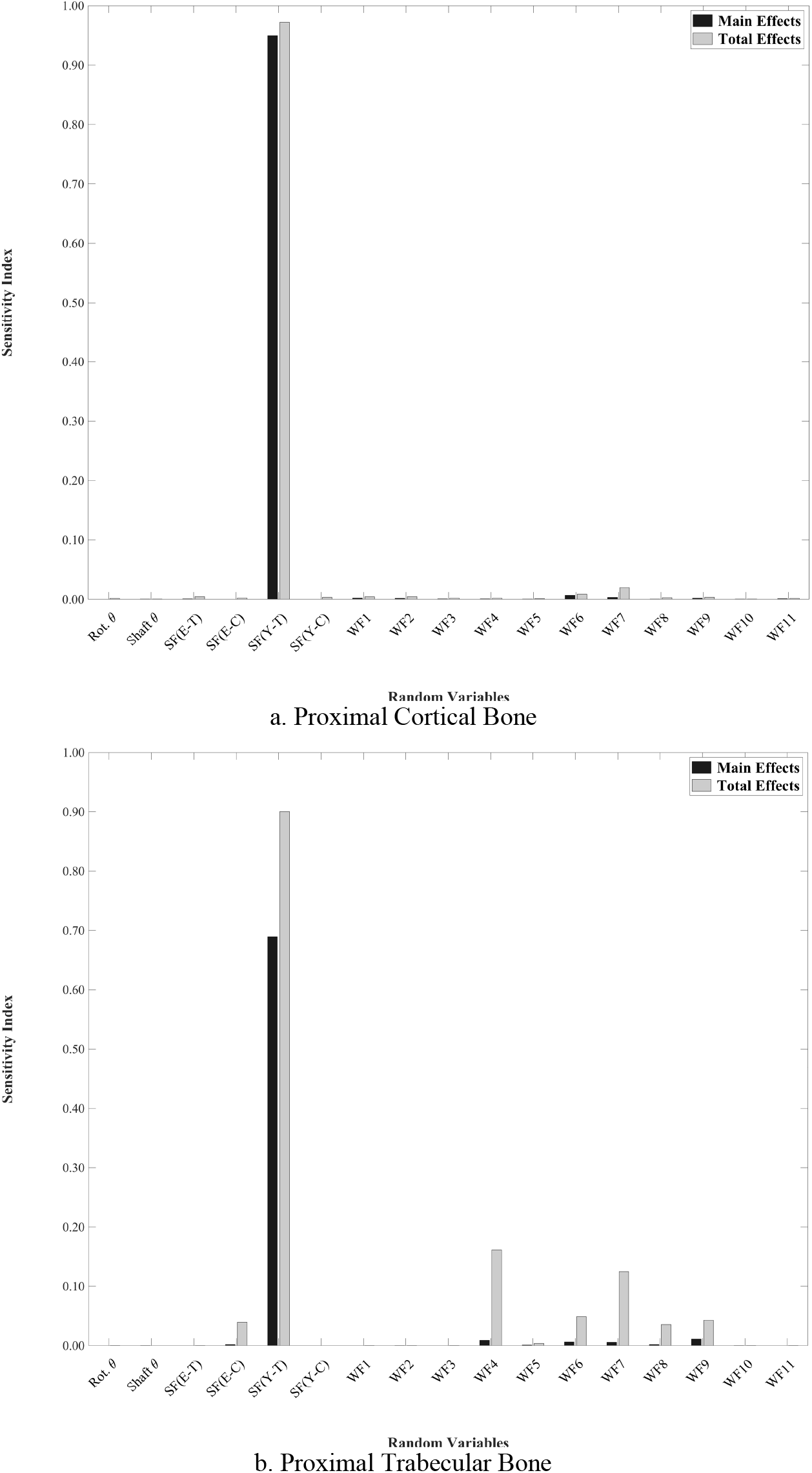
Association between random variables and stress response in the proximal femur under fall loading conditions (Rot. q = internal rotation angle, shaft q = shaft angle, SF(E-T) = trabecular modulus scale factor, SF(E-C) = cortical modulus scale factor, SF(Y-T) = trabecular yield strength scale factor, SF(Y-C) = cortical yield strength scale factor, WF = weighting factor)

**Figure 3:**
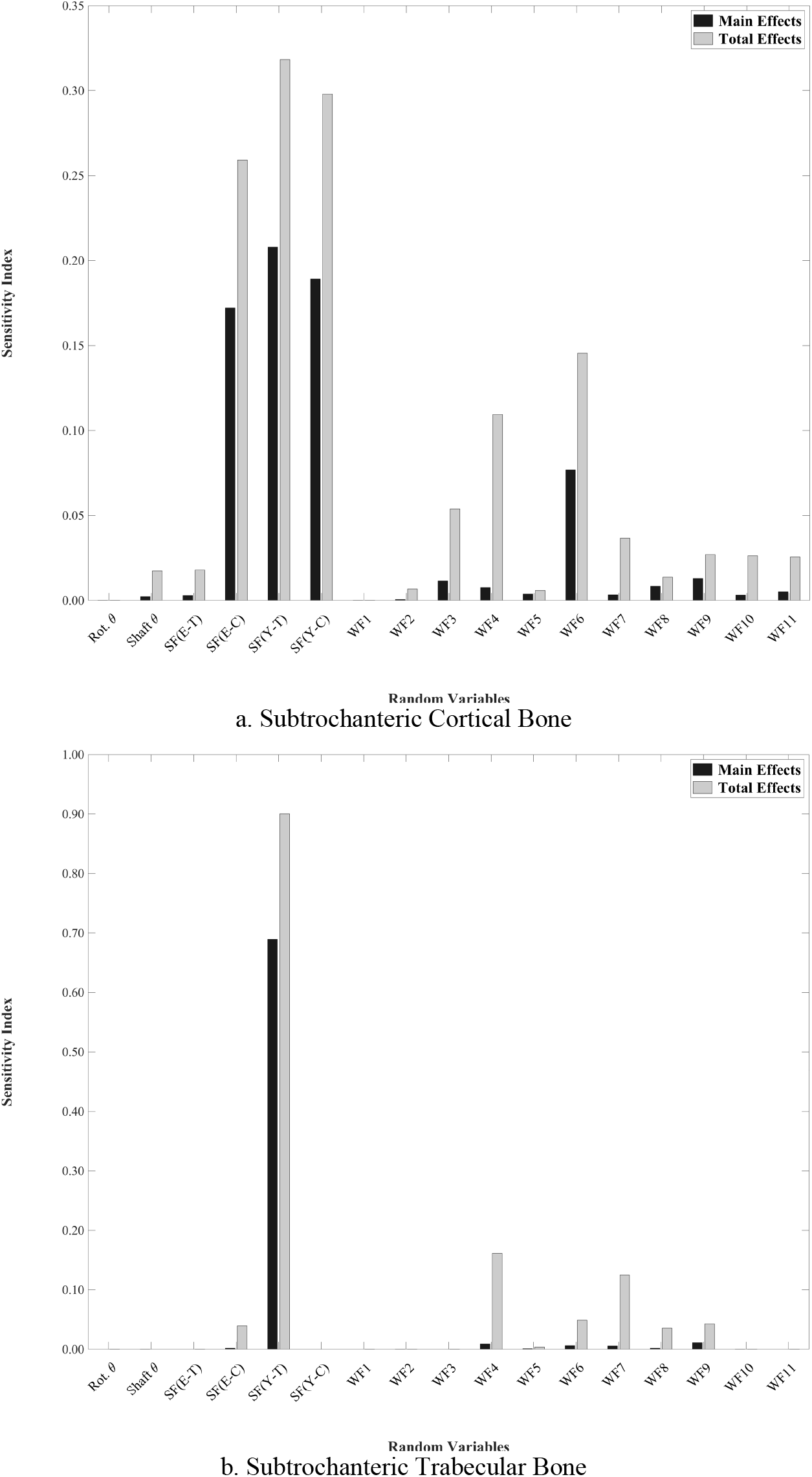
Association between random variables and stress response in the subtrochanteric femur under fall loading conditions (Rot. q = internal rotation angle, shaft q = shaft angle, SF(E-T) = trabecular modulus scale factor, SF(E-C) = cortical modulus scale factor, SF(Y-T) = trabecular yield strength scale factor, SF(Y-C) = cortical yield strength scale factor, WF = weighting factor)

Stress variation under simulated stance loading conditions was dependent on both the individual effects of loading, material properties, and femur structure variation. The stress distribution in the proximal femur was largely dependent on interaction between loading conditions and femur structure, although yield strength of the trabecular bone also contributed to proximal femur stress (Figure 4). As with fall loading conditions, the subtrochanteric stress distribution was dependent on material properties of the trabecular bone and femur structure, as well as interactions with variation in loading conditions (Figure 5).

**Figure 4:**
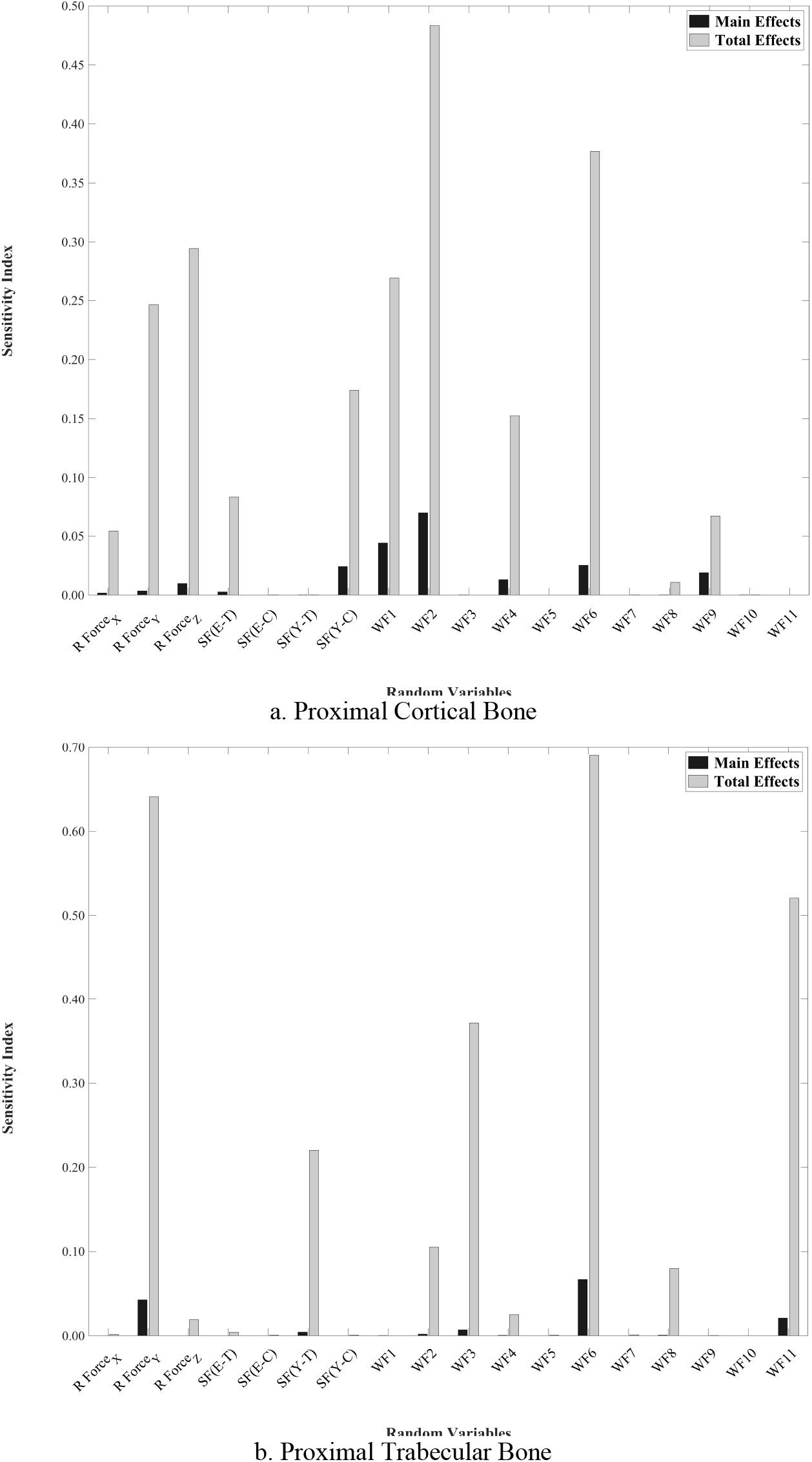
Association between random variables and stress response in the proximal femur under stance loading conditions (R Force_X,Y,Z_ = femoral resultant force components, SF(E-T) = trabecular modulus scale factor, SF(E-C) = cortical modulus scale factor, SF(Y-T) = trabecular yield strength scale factor, SF(Y-C) = cortical yield strength scale factor, WF = weighting factor)

**Figure 5:**
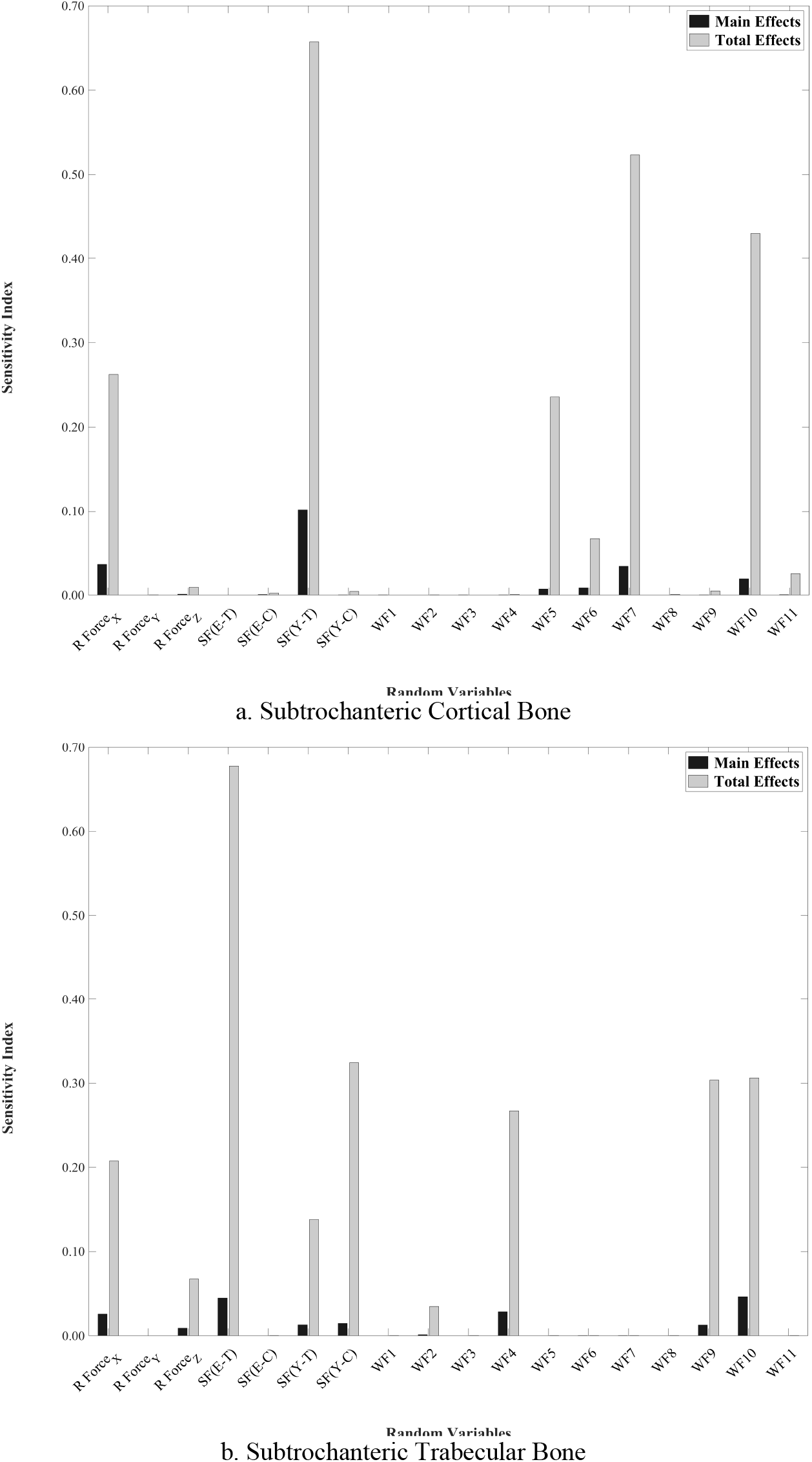
Association between random variables and stress response in the subtrochanteric femur under stance loading conditions (R Force_X,Y,Z_ = femoral resultant force components, SF(E-T) = trabecular modulus scale factor, SF(E-C) = cortical modulus scale factor, SF(Y-T) = trabecular yield strength scale factor, SF(Y-C) = cortical yield strength scale factor, WF = weighting factor)

### Associations between femur structure variation and weighting factors

Variations in typical geometric and densitometric measures of femur structure were associated with combinations of weighting factors and, conversely, each weighting factor described some variation in a large number of structural measures (Table 2). For instance, the first weighting factor described 81% of the variability in the set of 47 femurs and that variability was associated with changes in femur curvature, particularly sagittal curvature of the full femur, cortical thickness in the subtrochanteric and midshaft regions, proximal femur geometry, and the distribution of bone mineral density in the proximal femur. Although less variability was described by the additional weighting factors, weighting factors still described relatively large changes in femur structure (Table 2).

**Table 2:** Percent variation in femur structure descriptors associated with parametric femur weighting factors

| Femur Descriptors | Parametric Model Weighting Factors |  |  |  |  |  |  |  |  |  |  |
| --- | --- | --- | --- | --- | --- | --- | --- | --- | --- | --- | --- |
|  | WF1 | WF2 | WF3 | WF4 | WF5 | WF6 | WF7 | WF8 | WF9 | WF10 | WF11 |
| Coronal Curvature (Proximal) | -16 | -49 | -47 | - | - | 22 | -33 | -29 | - | 5 | -45 |
| Sagittal Curvature (Proximal) | - | 41 | 38 | 261 | - | 49 | 12 | -6 | 14 | - | 26 |
| Coronal Curvature (Midshaft) | -37 | 20 | -16 | -13 | 50 | 21 | -44 | -35 | -37 | -24 | 55 |
| Sagittal Curvature (Midshaft) | -69 | -58 | -57 | -50 | -57 | -47 | -55 | -69 | -52 | 223 | 22 |
| Coronal Curvature (Full) | -29 | 8 | -7 | 12 | 17 | - | -17 | -13 | -12 | - | 19 |
| Sagittal Curvature (Full) | 830 | -69 | -51 | -69 | -66 | 24 | -72 | 24 | -62 | - | -25 |
| Cortical Thickness (Subtroch. Mean) | -17 | 16 | - | -11 | -8 | 25 | - | -8 | -7 | -13 | 5 |
| Cortical Thickness (Subtroch. Min) | 7 | -13 | 17 | -5 | 20 | 104 | 10 | -38 | 54 | - | - |
| Cortical Thickness (Subtroch. Max) | -13 | 26 | 13 | - | -6 | 20 | - | - | -20 | -9 | 14 |
| Cortical Thickness (Midshaft Mean) | -18 | 22 | -5 | -8 | -16 | 18 | - | - | -12 | -15 | 5 |
| Cortical Thickness (Midshaft Min) | -9 | - | -11 | - | -57 | 20 | -22 | -31 | -16 | -38 | 7 |
| Cortical Thickness (Midshaft Max) | -17 | 23 | - | - | -9 | 20 | 7 | - | -6 | -9 | 11 |
| Neck Diameter | -14 | 9 | -11 | -11 | - | -6 | - | 7 | - | - | - |
| Neck Length | -50 | 87 | 43 | - | 20 | - | 38 | -11 | - | 11 | -34 |
| Neck Axis Length | -20 | 9 | - | -6 | - | - | - | - | - | - | - |
| Neck Axis - Shaft Angle | - | - | -60 | - | - | - | -57 | - | - | - | - |
| Femur Length | -18 | - | - | - | - | - | - | - | - | - | - |
| Mean Density (Neck, Cortical) | -13 | - | -15 | - | 13 | 9 | - | -9 | -8 | 7 | - |
| Mean Density (Neck, Trabecular) | -9 | - | -17 | - | - | 13 | - | - | - | 0 | -11 |
| Mean Density (Intertrochanteric, Cortical) | -26 | - | 26 | 25 | -8 | 21 | - | - | 45 | -21 | -23 |
| Mean Density (Intertrochanteric, Trabecular) | -22 | 6 | 68 | 53 | - | 33 | 25 | 13 | 63 | -18 | -23 |
| Mean Density (Subtrochanteric, Cortical) | - | 9 | - | 6 | -14 | 16 | - | 10 | 13 | -12 | - |
| Mean Density (Subtrochanteric, Trabecular) | - | -7 | - | - | - | 20 | -8 | - | - | - | -10 |
| Mean Density (Midshaft, Cortical) | - | 11 | - | - | - | - | - | - | - | - | - |
| Mean Density (Midshaft, Trabecular) | -12 | - | - | -17 | - | 18 | - | - | - | -7 | - |

Note: Structural variation is given as a percent change with respect to structural descriptors of the mean femur.

## DISCUSSION

Significant variability was present in the structural organization of a relatively small set of femurs, both in terms of femur morphometry, as well as in the associated spatial distribution of cortical bone thickness and bone density within the femur structure. Statistical shape and trait modeling is capable of efficiently describing variability in the complex structural organization of the femur and extends naturally to computational simulations allowing investigations of the effects of such variability on the loading response of the femur. In combination with parameters describing load condition and material property variation, variation in structural organization is associated with increases in tensile stress by region, which are likely associated with increased fracture risk.

Increased maximum principal stress was observed in both trabecular and cortical bone compartments of the upper femur, particularly in the intertrochanteric and subtrochanteric regions for models subjected to sideways fall and stance loading conditions. There were also a number of variation models where the tensile stress increases led to maximum principal stress values greater than corresponding tensile yield strength, values indicating that variation in parameters describing femur structure, material properties, and loading conditions are likely associated with increased fracture risk.

The multiple patterns of variation in descriptors of femur structure, loading conditions, and material properties that were associated with increased tensile stress values are complex and varied by region and within both trabecular and cortical bone compartments. Based on the results of computational simulations described here, it seems likely that even small variation in femur structure, loading, and material properties may lead to increased regional tensile stress and fracture risk. It also seems likely that shifts in the tendency towards typical proximal femur fracture and atypical femur fracture result from combinations of structural, loading, and material parameters.

In the development of computational models, there is inherent uncertainty in the model parameters specifying bone structure and structural organization, material properties of musculoskeletal tissue, and the loading conditions that might reasonably be experienced during low-energy falls or normal ambulatory activity. However, few studies in the literature investigate the effects of uncertainty in computational model definition, despite indications that results of musculoskeletal modeling simulations, even in the case of subject-specific models, are affected by these uncertainties [49]. Quantifying the effects of uncertainty on variation in model response by developing computational models within a probabilistic framework is a strength of the present study, importantly allowing the consideration of the effects of multiple input parameters, but also critically allowing for examination of the effects of interaction between structure, loading, and material behavior.

The results of this study indicate that there are multiple pathways and combinations of loading, material property, and femur structure variation that may result in increased fracture risk and that these pathways can lead to increased tensile stress and possible fracture, in any region of the femur under both overload conditions, such as with sideways fall loading, and stance loading. The repetitive nature of stance loading may lead to accumulation of fatigue damage within the bone, impairing bone condition and increasing susceptibility to fracture. In fact, it seems likely that damage accumulation resulting from cyclic tensile strains experienced during the stance phase of walking is associated with the location of atypical femur fractures [28]. Repetitive low-grade loading has commonly been associated with tibial stress fractures in military recruits [50, 51]; however, similar fatigue fractures have also occurred in the femoral neck of both military recruits (i.e., young adults) in basic training [52] and in the elderly [53]. Combinations of variation in femur structure, material properties, and loading conditions were all associated with variation in the femur stress distribution in stance loading. Accordingly, it is likely that some individuals may be predisposed to stress fractures and atypical femur fractures, particularly in the case of impaired material properties of bone.

Long-term bisphosphonate therapy to treat the reduced bone mineral density associated with osteopenia and osteoporosis has been associated with increased mineralization and embrittlement of cortical bone, leading to decreased ductility and postyield toughness [54]. The resulting inability of bone tissue to withstand normal, repetitive loading has been observed as increased tendency for initiation and accumulation of damage (e.g., microcracks) in bone [55]. Material property variation, particularly in bone ductility, has the predominant effect on subtrochanteric cortical bone under stance conditions in the present study and plays a major role in the stress distribution in subtrochanteric cortical bone under fall conditions.

Similarly, type 2 diabetes (T2D) has been shown to have a negative effect on the collagen matrix in trabecular bone, which results in decreased material properties, particularly in post-yield behavior [56]. Again, the yield behavior of trabecular bone has a strong association with subtrochanteric bone response, both in cortical and trabecular bone, in the present study. It seems clear that T2D is associated with higher fracture risk in fractures of the proximal femur, despite unchanged or increased volumetric bone mineral density compared to non-diabetics [57, 58]. However, the role (if any) of T2D in atypical femur fracture development is much less clear, despite inclusion as a risk factor in the current ASBMR Atypical Femur Fracture Taskforce report [21]. Giusti, et al. report a significant association between diabetes and low-energy subtrochanteric femur fractures [59]; however, in a retrospective study of closed femur fracture cases in south Texas, where the prevalance of T2D is over 25% in the elderly population, there was no significant difference in T2D incidence between those with subtrochanteric or midshaft femur fractures and those with intertrochanteric fractures [60]. The present study demonstrates that increased risk of either proximal femur or atypical femur fracture is associated with material property variation, as has been shown to occur with either long-term bisphosphonate use or T2D, but that femur structure also has a role in variation in the loading response. Taken as a whole, these findings indicate the critical importance of considering musculoskeletal trait covariation and variation in the loading environment in determining fracture risk and tendency towards typical or atypical femur fractures.

In the present study, loading direction was associated with variation in the stress distribution of subtrochanteric cortical bone. Similarly, an association was found between standing lower limb alignment and the location of fracture in atypical femur fractures in a retrospective review of ten patients with atypical femur fractures, including four patients with bilateral atypical femur fractures [22]. Pinilla, et al. showed that even slight variation in impact direction during a low-energy fall is associated with a significant variation in failure load of the proximal femur, as is also indicated in the stress variation in both cortical and trabecular in the present study [32].

The parametric femur model describes structural variation in a relatively small set of femurs from a relatively homogeneous sample of the population, as is often the case in studies involving postmortem human subjects. As such, only limited variation in femur structure was considered. For instance, femur curvature, particularly in the sagittal plane, is more pronounced in the Japanese population, regardless of age, gender, and femur length, than in the American White, African-American, and Asian American populations [61]. This study did not investigate the role of increased levels of variation in bone structure, as the intent was to investigate whether any variation in structure, material, and loading, either in isolation or as the effect of interaction with other variations would lead to variation in the magnitude and location of the femur stress response. Additionally, a relatively simple definition of material property assignment was employed here and was similar to that of other studies [62] [63] [64] [65]. More complicated models of material behavior and other response measures have been employed in the literature to investigate fracture risk [64, 66, 67]. However, the goal of the present study was to simply investigate the existence of associations between factors that are expected to be uncertain or to vary in the context of femur fracture risk and it is not clear that additional model complexity would affect the outcomes of the present study. Additionally, the mean response predicted using computational simulations of individual femurs under fall loading conditions was greater than that of experimental femurs. It is not clear whether these differences in response were associated with structural variations between the groups of femurs, but as the range of responses were similar and relative behavior between femurs was considered in the context of the effects of variation in the present study, it is not likely that the results of the present study are affected.

## CONCLUSIONS

Variation in femur structure, material behavior, and loading conditions were associated, both individually and in combination with variation in the structural response of the femur under fall-type and stance loading. Multiple patterns of variation in structural descriptors were associated with increases in regional stress in both the proximal and subtrochanteric regions and in cortical and trabecular bone, suggesting that there are multiple pathways that lead to increased fracture risk, either in the case of typical proximal femur fractures or atypical femur fractures. Consideration of the uncertainty in model definition is a critical, but seldom-employed, approach to advances in the etiology of bone fractures. It seems clear that bone fracture and the location of fracture is not the result of a limited set of musculoskeletal traits or loading condition variables, but, rather, is the result of complex interaction between all of these factors.

## Acknowledgment

This work was sponsored in part by Merck Investigator Studies Program Award #55715.

